# Data-centric AI approach for automated wildflower monitoring

**DOI:** 10.1101/2024.04.18.590040

**Authors:** Gerard Schouten, Bas S.H.T. Michielsen, Barbara Gravendeel

## Abstract

Both researchers and policy makers are in need of standards and tools that help understanding and assessing natural capital. Wildflowers are a major component of our natural capital; they play an essential role in ecosystems, improve soil health, supply food and medicines, and curb climate change. In this paper, we present the Eindhoven Wildflower Dataset (EWD) as well as a PyTorch object detection model that is able to *identify* and *count* wildflowers. EWD, collected over two entire flowering seasons and expert annotated, contains 2002 top-view images of flowering plants captured ‘in the wild’ in five different landscape types (roadsides, urban green spaces, cropland, weed-rich grassland, marshland). It holds a total of 65571 annotations for 160 species belonging to 31 different families of flowering plants and serves as a reference dataset for automating wildflower monitoring. To ensure consistent annotations, we define specific floral count units (largely based on inflorescences) and provide extensive annotation guidelines. With a 0.82 mAP (@IoU > 0.50) score the presented baseline model, trained with a balanced subset of EWD, is to the best of our knowledge superior in its class. Our approach empowers automated quantification of wildflower richness and abundance and encourages the development of standards for AI-based wildflower monitoring. The annotated EWD dataset is publicly available on the DataverseNL research data repository, and the code to train and run the baseline model is supplied as supplementary material.

## 1. Introduction

Wildflowers, or angiosperms, are plants producing flowers that, once pollinated, develop into seed-bearing fruits. They are a crucial part of our ecosystem. In the history of life, they are relative newcomers. The first flowering plants emerged about 130 million years ago – approximately 70 million years after the first appearance of mammals – following a period of rapid evolution [1, 2]. At present, there are more than 300,000 species of flowering plants worldwide [3]; they outnumber all other plants combined and are the dominant vegetation on land [4]. Flowers exhibit a large diversity in color, shape and texture to attract pollinators. Besides playing crucial roles in natural ecosystems, plants provide a great variety of essential services to human life. Root systems of plants improve soil health, purify water, and prevent erosion. In addition to their aesthetic and cultural value, plants strongly regulate daily temperature variations in built environments [5] and provide building materials, food as well as a plethora of medicines [6, 7]. Moreover, the photosynthesis process of plants mitigates climate change by capturing carbon dioxide.

An imperative condition for sustainable ecosystem services, as the ones mentioned above, is a diverse and resilient web-of-life [8]. Today, ecosystem services are degrading because of biodiversity loss [9]. To better understand ecosystems and maintain their services it is essential to have automated large-scale monitoring programs to assess biodiversity trends reliably and consistently over long periods of time [10–12].

Current practice for wildflower monitoring includes systematic surveys that rely on manual counts by professionals and nature enthusiasts complemented with data collection supported by citizen-science platforms. These routines have limitations hampering large-scale monitoring. Manual counting is arduous, error prone, and labor-intensive. Species identification – although supported nowadays by well-designed AI-enabled smartphone apps, such as Pl@ntNet [13] or PlantSnap [14] – is still challenging for the general public. The situation is further exacerbated by the fact that taxonomic knowledge is becoming increasingly scarce [15, 16]. Another drawback, especially for the more ad-hoc data collection in open citizen-science platforms, is biased data [17]. Orchids or other ‘iconic’ flowering plant families are overrepresented, while less sexy ones are underrepresented.

Given these issues, wildflower monitoring would be greatly helped by the development of AI solutions that can perform reliable automated wildflower counts from images of patches of land. The smartphone apps mentioned earlier cannot do this. They are driven by AI models that have been trained on photographic close-ups of flowers, and only solve a *classification* problem, implicitly assuming that each new input contains only one item. Classifying and localizing multiple wildflowers in overview images is a different, more complex challenge that requires an *object detection* approach [18]. Although visual object detection is advancing rapidly in other domains (e.g. autonomous driving and healthcare), there are to our knowledge currently next to no solutions in the domain of wildflower monitoring.

It is our aim to make a first step towards creating an AI solution that can carry out reliable automated counts from overview images containing multiple wildflowers of various species. This challenge involves two major elements: (i) the creation of an annotated dataset of such overview images, and (ii) the creation of an object detection model trained on the annotated dataset.

In this paper, we present:

1. The Eindhoven Wildflower Dataset (EWD). This is an expert-annotated reference dataset of over 2000 bird’s-eye view high-resolution wildflower images, captured in five different Dutch landscape types – roadsides, urban green spaces, extensively farmed cropland, weed-rich grassland, and marshland. We limited ourselves to herbs and collected images in two entire flowering seasons.
2. A state-of-the-art object detection model trained on EDW. Our model is based on off-the-shelf open-source software and serves as a baseline. It is intended as a steppingstone for new innovative modelling efforts in the domain of automated wildflower monitoring.

Many deep learning papers aim at improving predictive models or creating novel, more efficient algorithms. Contrary to this algorithm-centric perspective, we adopt a data-centric AI approach. This refers to a recent shift visible in the literature from focusing on the performance of models to the quality of the underlying data used to train and evaluate these models [19]. This is a fundamental change, where data is viewed as a profound asset, on par with model code. To obtain high-quality data for wildflower monitoring, we formulate explicit annotation guidelines [20] and introduce floral count units, compatible with botanical nomenclature. Moreover, before training the model we created a balanced subset of EWD and tiled the images to preserve resolution.

## 2. Background and related work

### 2.1 Datasets as benchmarks

The role of image-based reference datasets varies from supporting the famous general-purpose computer vision competitions, such as PASCAL VOC [21], COCO [22], ImageNet Large Scale Visual Recognition Challenge [23], and Open Images V4 [24] to benchmarks for domain-specific quests. For instance, in the medical domain open datasets with skin lesion images played a major role in creating models that can diagnose malicious melanomas [25–27] or, more recently, public CT and X-ray chest data significantly accelerated the development of clinical apps that are able to estimate COVID-19 infection status [28, 29]. Similarly, the Audi A2D2 dataset [30] pushes forward the research field of autonomous driving. Even synthetic data is utilized as reference data, e.g., for simulating driving scenarios with fast context changes, such as deteriorating weather conditions [31].

In short, the proliferation of annotated image-based public datasets in the scientific literature fast-tracks the creation of novel model architectures [32] and improvement of evaluation metrics, e.g. [21, 33]. It allows the scientific community to compare computer vision algorithms and pushes the field towards increasingly complex and challenging problems. Furthermore, they trigger more strict annotation protocols to raise data quality [20].

### 2.2 Public image-based flower datasets

Table 1 lists the most important public datasets with flower images taken ‘in the wild’ that target either the species identification or monitoring use case. The top six datasets in Table 1 include only close-up images of individual wildflowers. The Kaggle dataset with only five very distinctive wildflower species mainly serves an educational purpose. It contains low resolution images scraped from Flickr and Google Images. The Oxford 17 dataset [34] has also been acquired by searching the web; it contains 80 images for each of the 17 flowering plant species included in this study from the UK; Oxford 102 [35] holds, as the name indicates, 102 plant taxa from the UK represented by 40 to 258 images per class. It is more challenging due to small inter-class variances and large intra-class variances. The Jena 30 dataset [36] consists of iPhone6 images of 30 wildflower species (11 to 70 images per class) and was collected during an entire flowering season in semi-arid grasslands around Jena in Germany. The PlantCLEF challenge [37] targets plant identification at a global scale. It is a fast-growing database currently holding 2,9 million images of about 80.000 species. It contains images of both flowers and leaves. The train data has two types of classification labels: either ‘trusted’ (validated) or ‘web’ (might suffer from species identification errors). The holdout test set consists of tens of thousands of pictures verified by experts. HDF100 extends flower identification into the hyperspectral domain [38].

**Table 1:** Public flower datasets with images taken ‘in the wild’.

| **Dataset (year)** | **# images \| type (# ann.)** | **# species** | **Purpose** | **Computer vision task** | **Annotation guidelines** |
| --- | --- | --- | --- | --- | --- |
| Kaggle Flower (--) | 4242 \| RGB | 5 | Educational challenge | Classification | No |
| Oxford 17 (2006) | 1360 \| RGB | 17 | Species identification | Classification | No |
| Oxford 102 (2008) | 8189 \| RGB | 102 | Species identification | Classification | No |
| Jena 30 (2017) | 1479 \| RGB | 30 | Species identification | Classification | No |
| PlantCLEF (2022) | 2,9 M \| RGB | 80K | Species identification | Classification | No |
| HFD100 (2022) | 10700 \| Hyperspectral | 100 | Species identification | Classification | No |
| Nectar (2019) | 1997 \| RGB (25K) | 25 | Wildflower monitoring, nectar index estimation | Object detection | No |
| **EWD (ours) (2024)** | **2002 \| RGB (65.6K)** | **160** | **Wildflower monitoring** | **Object detection** | **Yes** |

The bottom two rows in Table 1 refer to datasets containing top-view images with tagged bounding boxes (BB) around wildflowers. Both support the monitoring use case, i.e., the use of object detection models that can *classify* and *localize*. The Nectar dataset [39] consists of nearly 2000 Canon Powershot G10 (14.7 Mpixels) images of flowering plant species collected in one flowering season in one habitat type, viz. weed-rich grasslands in the UK. All 25 species are easy to outline and have predominantly simple inflorescences, i.e., solitary flowers or flower heads. The aim of the Hicks et al. study [39] was to estimate nectar sugar mass as an indicator for effectiveness of nature-inclusive farming. In other words, these authors used wildflower monitoring as a means to an end. In our study, wildflower monitoring with computer vision is the main topic. Although the EWD dataset has a comparable number of Canon EOS Mark IV (30.4 Mpixel) images, it holds a significantly larger number of flowering plant species. The EWD images were collected over two entire flowering seasons in five different habitat types around the city of Eindhoven in the Netherlands. Even more importantly, what makes EWD novel and unique, is that the annotated images adhere to strict guidelines and that well-defined floral count units are used, also for more complex inflorescences. In short, EWD complies with the data-centric AI principle to boost data quality.

### 2.3 Challenges and solutions for automated wildflower monitoring

Automated wildflower monitoring has its own unique computer vision challenges. Backgrounds can be very cluttered, and flowers ‘in the wild’ are often occluded or damaged. Moreover, flowers develop from bud to blooming stage and subsequently whither or change into fruits, i.e., they are not stable in size, shape and color but change during their life cycle. And above all, wildflower identification is a large vocabulary problem with few majority classes and many minority classes [40]. This requires a classifier that can handle class imbalance and that supports fine-grained class differentiation, i.e. small inter-class variances (related flowering species might be quite similar to one another) as well as large intra-class variances (individuals of the same species may vary considerably in their morphology). Fig 1 illustrates some of these challenges with EWD images.

**Fig 1:**
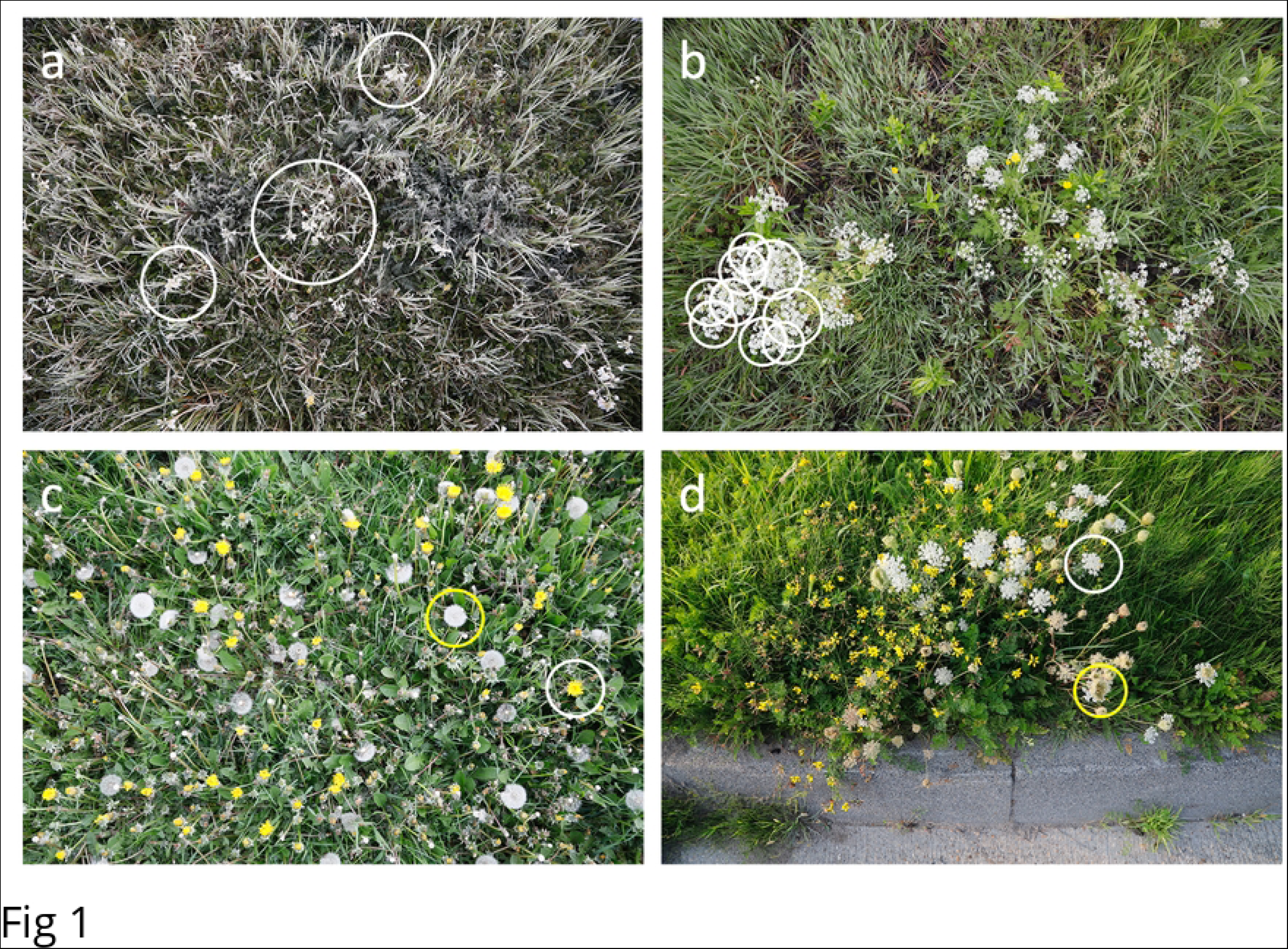
Computer vision challenges for top-view wildflower images taken ‘in the wild’. The panels show from top left to bottom right: a) *Cardamine pratensis* is difficult to detect on a cluttered background with a touch of frost, b) overlapping umbels of *Anthriscus sylvestris* are hard to count, c) wildflowers change over time, *Taraxacam officinale* in blooming (white circle) and fruiting stage (yellow circle), and d) *Daucus carota* in blooming (white circle) and fruiting stage (yellow circle).

#### 2.3.1 Classification

Let us first focus on the ‘easier’ *classification* problem. Over the years various computer vision algorithms have been proposed to deal with these challenges. A straightforward method to identify wildflowers in images is explicitly coding morphological features (colour, texture, shape) with handcrafted image processing filters [41, 42]. More elaborate approaches compute local image descriptors [43], such as *scale-invariant feature transform* [44] or *histogram of gradients* [45], followed by a machine learning classifier. The descriptor or early learn-from-data-implementations resulted in significantly higher precision and recall rates compared to the algorithms without machine learning [35, 36]. An extensive survey of wildflower identification with traditional image processing as well as the descriptor-classifier approach is given by Wäldchen et al. [46].

Since the landmark paper of Krizhevsky et al. [47] – in which an 8-layer convolutional neural network (CNN) was presented that won the ILSVRC challenge – and the subsequent designs of even more powerful CNN architectures [48–50], wildflower identification (along with many other computer vision problems) entered the realm of deep learning. Training a CNN with generalization capabilities requires a large, labelled dataset with intra-class diversity. A popular and efficient deep learning approach is transfer learning [51], in which a large convolutional neural network with pretrained weights is used as a visual encoder followed by a shallow classifier network that is trained to discriminate (or learn) several domain-specific object categories, such as flowering plant species. Currently, deep learning is state-of-the-art for flower identification. Using this technique, Wäldchen et al. [52] report accuracy levels of approximately 95% for the Oxford 102 benchmark dataset (containing 102 species, see Table 1), thereby outperforming the best descriptor-classifier result.

#### 2.3.2 Object detection

Rather than identifying flowers (from isolated or cropped-out close-ups), we aim to apply an *object detection* algorithm that can reliably identify and count wildflowers from overview images. Object detection seeks to locate and recognize object instances (from a large number of predefined classes) in natural images. For recent surveys of the developments in object detection, see [18, 53]. Adopting object detection for to wildflower monitoring adds a further challenge to the ones mentioned above. Whereas a ‘simple’ classifier answers the question “*What object is in the image?”*, object detection models answer the question “*What objects are where in the image?*”. Answering the latter question supports the monitoring use case, i.e., identifying wildflower instances and counting them.

Object detection models are trained with images annotated with tagged BBs around objects of interest. The annotations can be formalized as set {(*b_i_*^*g*^, *c_i_*^*g*^)}, where *b_i_*^*g*^ denotes the ground-truth BB of object *i* in the image, and *c_i_*^*g*^ the ground-truth class label. In inference mode, an object detection model takes an image as input and the standard output is a set of object proposals, each with an ‘objectness’ score. This results in the set {(*b*_*j*_,*c*_*j*_, *p*_*j*_)}, where *b*_*j*_ denotes the predicted BB, *c*_*j*_ the predicted category, and *p*_*j*_ the corresponding confidence score (indicating the likelihood of membership to a set of objects *vs.* background).

The metric to evaluate object detection models is mean averaged precision (mAP). It is a figure of merit between 0 and 1 expressing the precision of an object detection model (a high score refers to a more accurate model). It is computed from the test set by applying the following steps.

1. *Determine true positive (TP) and false positive (FP) object proposals*. When a prediction for a certain annotated object (*b_i_*^*g*^, *c_i_*^*g*^) satisfies the following three conditions: i) the predicted and ground-truth BBs, *b*_*j*_and *b_i_*^*g*^ respectively, have a sufficient degree of overlap, ii) the predicted label *c*_*j*_ is correct, and iii) the confidence score *p*_*j*_is higher than a preset threshold, it is considered as a TP. Otherwise, it is considered as a FP. The ‘sufficient degree’ of overlap condition is operationalized with the area of intersection over union for the predicted and ground-truth BB (i.e., (*b*_*j*_ ∩ *b_i_*^*g*^) / (*b*_*j*_ ∪ *b_i_*^*g*^), abbreviated IoU. An IoU > 0.5 is generally considered as a good prediction if precise localization is not a priority [21]. In the research presented in this paper we follow this common practice.
2. *Compute average precision (AP) per class*. Based on the TPs and FPs, precision-recall pairs can be constructed by varying the confidence score threshold. From the obtained precision-recall pairs (allowing precision to be regarded as a downwards-sloping function of recall), the average precision (AP) per class can be derived from the area under the curve [23].
3. *Average the AP scores*. Finally, mAP is calculated by averaging the per class AP scores.

In practice, an object detection model will now and then predict multiple BBs that sufficiently overlap and have matching class labels with above-threshold confidence scores. In this case the prediction with the highest IoU is the TP, duplicates are designated as FPs.

There are two categories of object detection models: Two-stage region-based CNNs (RCNNs) [54–56], and one-stage detectors such as single shot detectors (SSDs) and the YOLO (You Only Look Once) algorithm family [57, 58]. RCNNs are in general more accurate, whereas one-stage models are known for their near real-time performance in inference mode [53]. Given the goal of this study, speed is of minor importance, so we focus on the RCNN models. In RCNN models, first of all category-independent region proposals are generated from an image with a so-called Region Proposal Network, next CNN-like features are extracted from these regions, and then category-specific classifiers are used to determine the category labels of the proposals.

So far, object detection has been explored sparsely for automatic wildflower monitoring. Ärje et al. [59] applied it to record plant phenology from time-lapse images; they targeted only one flowering plant species on Greenland: *Dryas integrifolia*. Hicks et al. [39], used it to derive a nectar sugar mass index from detected floral resources. And Gallman et al. [60] adopted this AI technology along with drone-based image acquisition and the construction of georeferenced orthomosaics for detecting wildflowers in mountainous areas. The proposed solutions of Hicks et al. [39] and Gallman et al. [60] can identify and count 25 common wildflowers. Note that the Gallman et al. [60] dataset is not open source, so it is not included in Table 1.

## 3. Eindhoven Wildflower Dataset

### 3.1 Collecting the EWD images

All 2002 EWD images encompass untrampled soil area of approximately 1m^2^ and were taken (near-)vertically downward from a height in the range of 1.5 – 1.9m. Most images (91%) were collected near the city of Eindhoven, the Netherlands. Data collection is limited to non-woody herbaceous flowering plants. Maximum allowed vegetation height was 1m. Images were acquired with 24-105mm/f4 lens mounted to a full-frame SLR camera (Canon Mark IV, 30.4 Mpixel). Exposure strategy: Fix ISO-setting at 800, maintain a shutter time faster than 1/50s to avoid motion blur caused by camera movement, and maximize aperture setting to create a large depth of focus.

EWD images were collected over two flowering seasons under various weather conditions in five different landscape types: i) roadsides (including dikes, bike trails, and railway borders; 832 images), ii) urban green spaces (parks, playgrounds, allotments; 237 images), iii) extensively farmed croplands (53 images), iv) weed-rich grasslands (457 images), v) marshland (including pools, fens, canals, and ditches; 423 images). The specific locations were selected not so much to create a dataset that is representative for the areas’ ecologies, but instead to mitigate the long-tailed species distribution one normally finds in biological data. In other words, using our knowledge of local landscapes, we aimed to include sufficient variation in the dataset, so that the model would have enough examples of a wide range of wildflowers to train on.

During the field work a log with a species list per image was recorded. Wildflowers were identified in the field using the 24^th^ edition of the Heukels’ Flora [61] supported by the ObsIdentify app [62]; the field worker (first author) has over 35 years of experience with botanizing. As said, the sampling strategy used can be summarized as a wisely chosen orchestration that ensures data collection with a fair number of wildflower species as well as a reasonably balanced number of instances per species. Contrary to Hicks et al. [39] we did not avoid human elements in the images, such as pavement, litter, sewer gullies, etc. The abovementioned measures guarantee a rich reference dataset for developing AI models that support real-life wildflower monitoring.

To prevent disclosure of exact locations of rare and red list wildflowers the metadata is removed from the collected images. No permits were required to collect the field data, all visited terrains (roadsides, urban parks, cropland, grassland, marshland) were open to the public and freely accessible.

### 3.2 Inflorescences and floral count units

All flowers are part of an inflorescence. Inflorescences vary in type among different plant families. An inflorescence type encodes both the branching pattern and the order in which its flowers open [63–65]. In this paper we adopt the formal, internationally agreed botanical inflorescence types. The inflorescences encountered in EWD are visualized in Fig 2. Some are relatively simple and form easy-to-outline structures (solitary flowers, spikes, and capitula or flower heads), others are more complex (dichasium, raceme, verticillaster, cyme, umbel, corymb, thyrse, panicle). For most wildflowers, inflorescences are excellent proxies for counting individual plants, as indicated by the EWD examples in Fig 3. By adopting inflorescences as floral count units (FCUs) we embody the building plan of wildflowers into the annotations. However, for a few species the standard inflorescences are impractical or ambiguous as floral count units in top-view relevées. For instance, it might be difficult or even impossible to outline the entire inflorescence, e.g., because the number of flowers varies too much (see Fig 4a), or it contains multiple flowers that typically bloom one at the time (see Fig 4b). In these circumstances we typically annotate single flowers. In case of nested inflorescences, the chosen FCU depends on how dense the secondary inflorescences are packed. As an example, for *Tanacetum vulgare* (corymb of capitula) the largest inflorescence type (corymb) is taken as FCU (see Fig 4c). One the other hand, for *Centaurea cyanus* (panicle of capitula) it is more convenient to take the individual flower heads as FCU (see Fig 4d). An important constraint for defining FCUs is to be consistent for look-alike species. For example: Since the inflorescence and the FCU of *Bellis perennis* is a capitulum, a flower head should also be used for other EWD wildflowers that have white ray florets and yellowish disc florets, like *Leucanthemum vulgare*, *Matricaria chamomilla*, and *Erigeron annuus*. The complete list of FCUs that we propose for each EWD species can be found in S1 Appendix. For 1 out of 5 flowering plant species the defined FCU does not have a one-to-one correspondence to its botanical inflorescence type.

**Fig 2:**
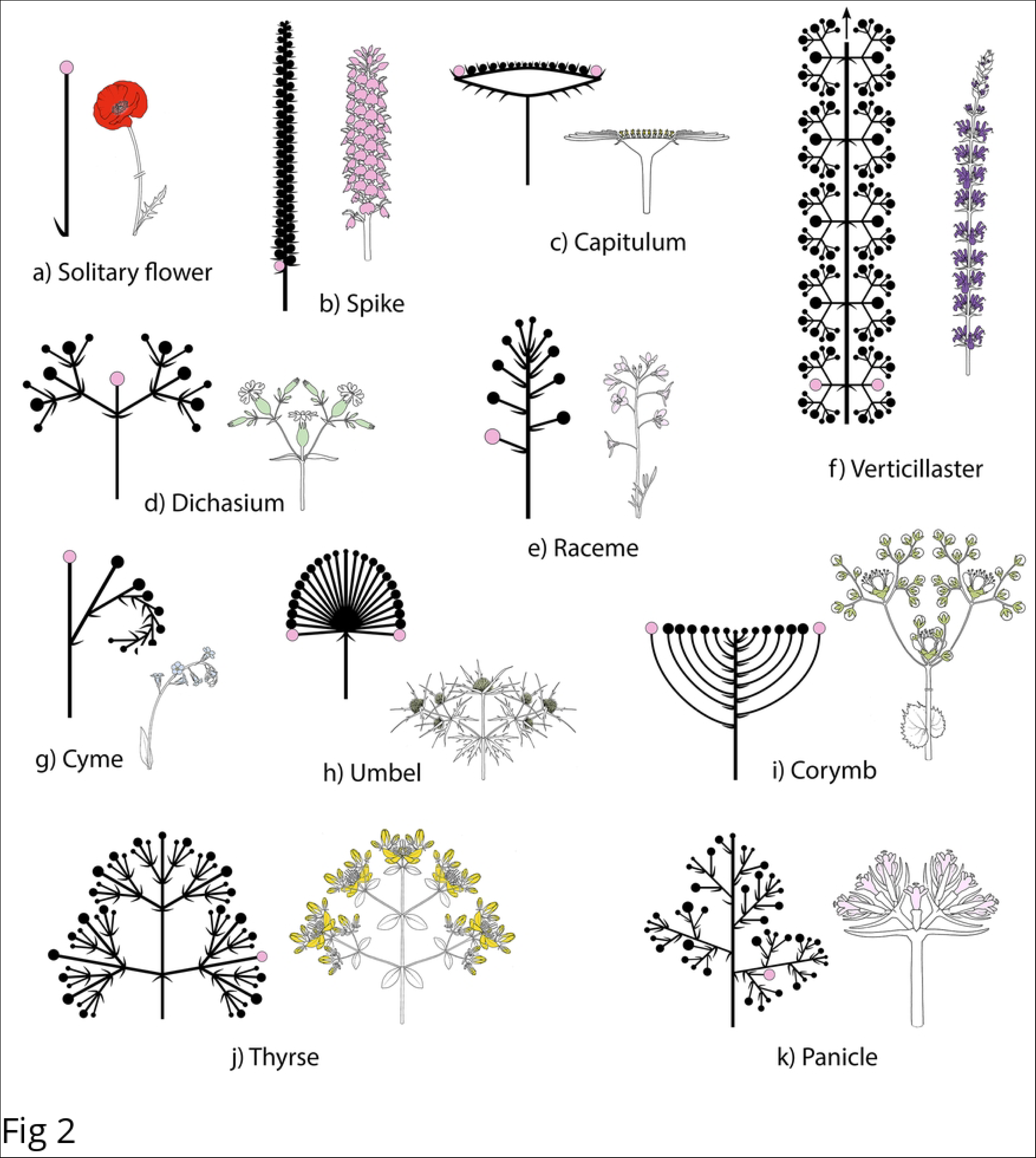
Inflorescences present in EWD. Schematic and example species. a) solitary flower – *Papaver rhoeas*, b) spike – *Dactylorhiza praetermissa*, c) capitulum – *Bellis perennis*, d) dichasium – *Silene latifolia*, e) raceme – *Cardamine pratensis*, f) verticillaster – *Salvia nemorosa*, g) cyme – *Myosotis arvensis*, h) umbel – *Erynchium campestre*, i) corymb – *Filipendula ulmaria*, j) thyrse – *Hypericum perforatum*, k) panicle – *Valeriana dioica*

**Fig 3:**
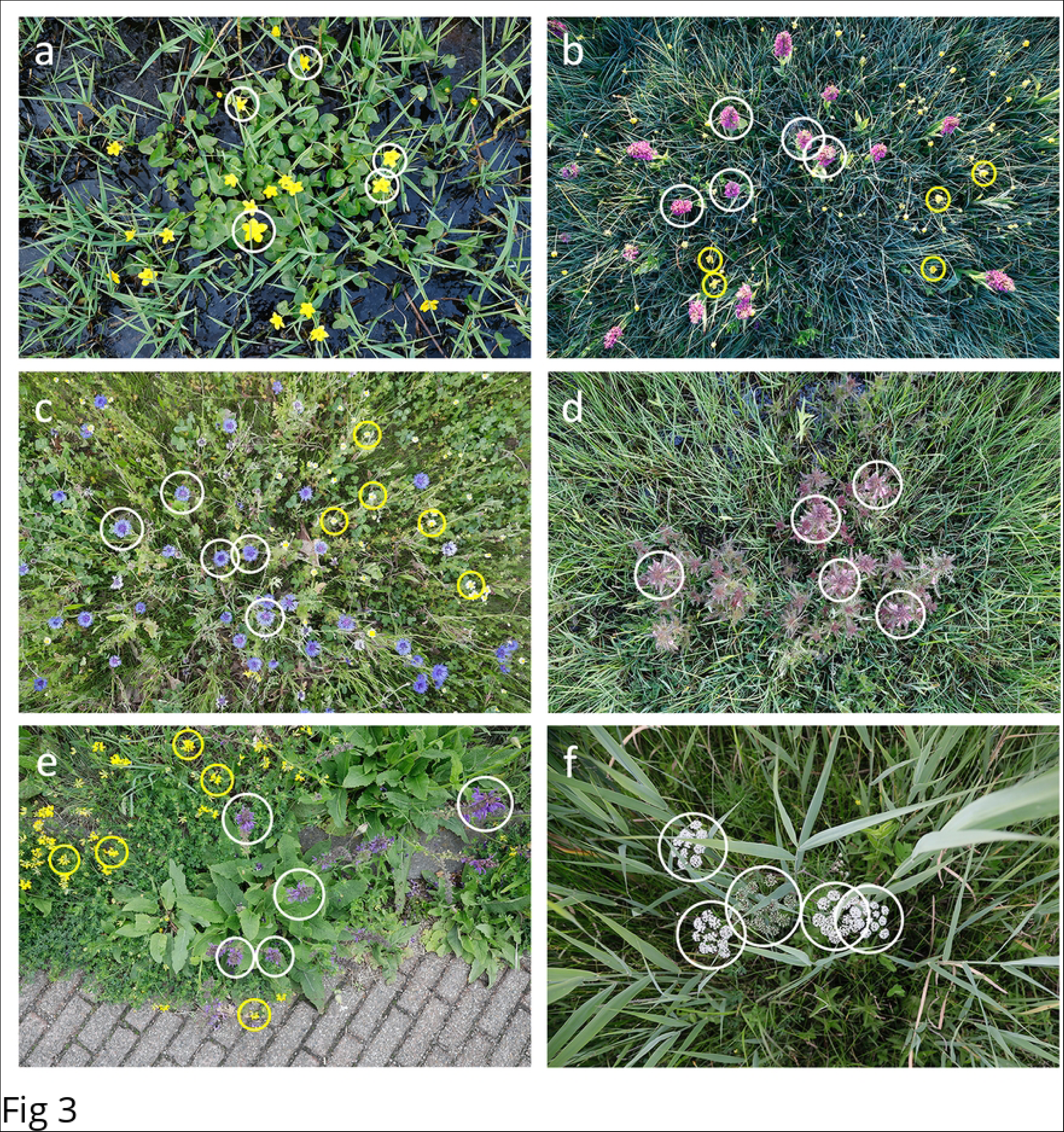
Examples of floral count units (FCUs) that match standard botanical inflorescence types. From top left to bottom right: a) FCU of *Papaver rhoeas* is a solitary flower, b) FCU of *Dactylorhiza praetermissa* is a spike, c) FCU of *Cirsium dissectum* is a capitulum (flower head), d) FCU of *Pedicularis palustris* is a raceme, e) FCU of *Salvia pratensis* (white circle) is a verticillaster, FCU of *Lotus corniculatus* (yellow circle) is a cyme, and f) FCU of *Cicuta virosa* is a compound umbel (typically containing 10 umbellets, each with up to 100 flowers). In each image only five FCUs are marked per species.

**Fig 4:**
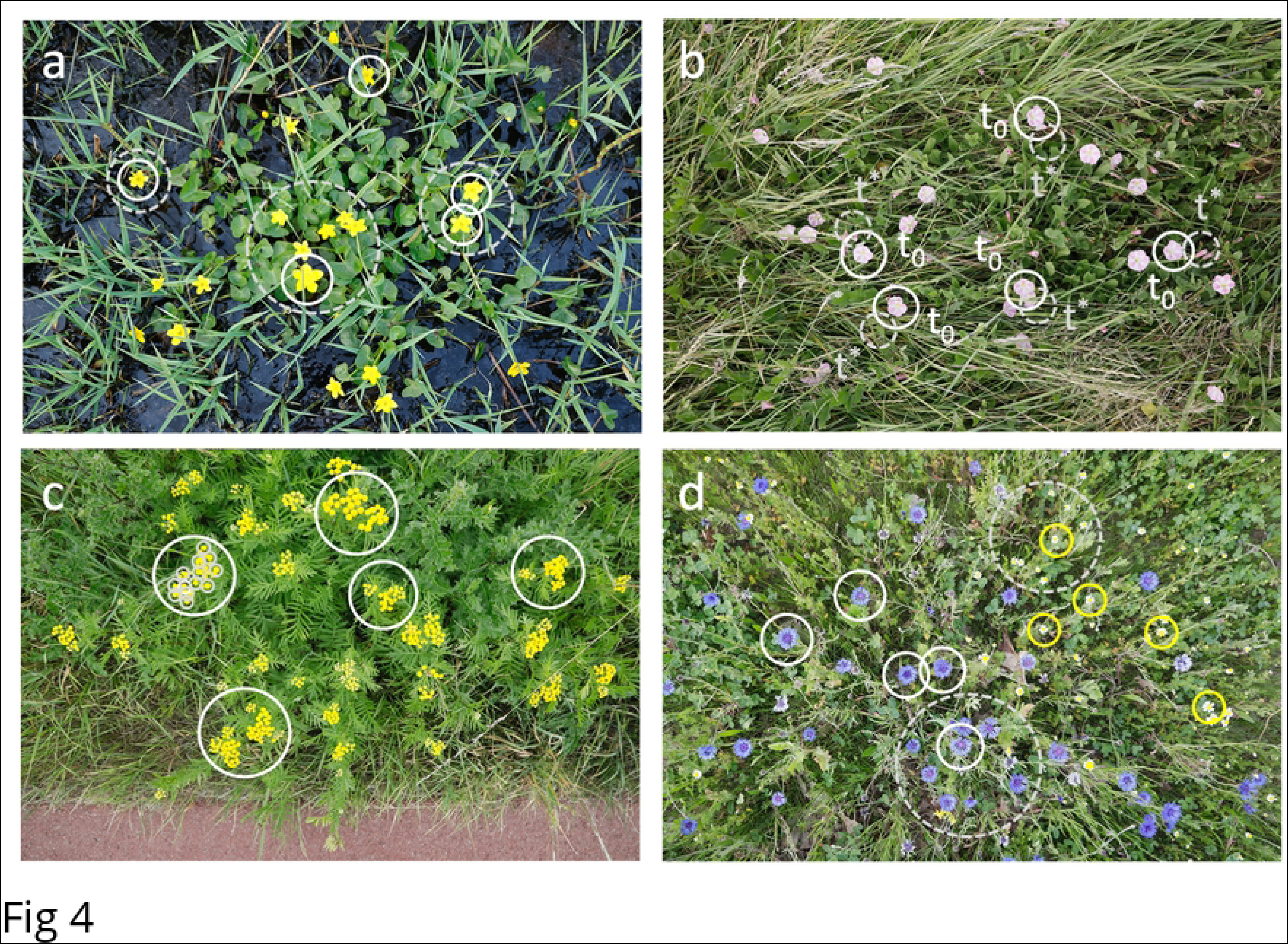
Examples of FCUs that do not match standard botanical inflorescence types. From top left to bottom right: a) The inflorescence of *Caltha palustris* is a cyme holding a variable number of 1 to 7 flowers. Since it is often impossible to outline these cymes (dashed grey circles are indicative) we therefore propose to use single flowers (white circles) as FCU. b) The stalked cyme of *Convolvulus arvensis* supports 1 to 2 flowers that bloom one after the other. White circles mark flowers that currently (*t_0_*) bloom; the adjacent grey dashed circles indicate possible future or past positions (*t\**) where flowers of the same cyme will appear (or disappeared). In this case we also propose to use single flowers as FCU. Right panel: c) The inflorescence of *Tanacetum vulgare* is a corymb of capitula. For this nested inflorescence it is convenient to use the flat-topped corymbs as FCU (large white circles) and not the individual flower heads (small grey dashed circles). d) Just the opposite; the inflorescence of *Centaurea cyanus* (white circle) and *Matricaria chamomilla* (yellow circle) is a panicle of capitula. It both cases it is impossible to outline the panicle (dashed grey circles are indicative) and we propose to use the individual flower heads (white circles) as FCU. In each image only five FCUs are marked per species.

Furthermore, each species-specific FCU is supplemented with two conversion factors obtained from manual inspection of digitized herbarium sheets collected in the Netherlands (near the city of Eindhoven) and registered as a collection item at the Naturalis Biodiversity Center. The first conversion factor represents the average number of flowers per FCU, and the second conversion factor represents the average number of FCUs per plant. These mappings extend FCU counts, as predicted by an object detection model, to other use cases such the assessment of nectar index, pollen density, winter food supply for birds, etc. and support demographic studies of plant populations.

In summary, what we register in each relevée are FCUs, defined for every species based on mostly practical considerations of countability. The FCUs can subsequently be converted into number of flowers or number of plants, using conversion tables.

### 3.3 Adding annotations to EWD

Adding annotations to real-world images is labour intensive and by nature a subjective process [66]. We took several precautions to objectify this process as much as possible. First, EWD images were annotated at species level by a plant expert making use of the field notes and respecting the species-specific FCUs (usually inflorescences) as listed in S1 Appendix. A few species that are visually indistinguishable in the field are merged: i) *Ranunculus acris* and *Ranunculus repens* are annotated as *Buttercup* (aggregate)*, ii) *Matricaria chamomilla* and *Matricaria maritima* are annotated as *Chamomile*\* *(aggregate),* and iii) unidentified yellow composites with only ray florets are marked as *Yellow Composite* (aggregate)*. Secondly, a 4- eyes principle was applied: Every annotation is double-checked.

To further materialize the annotations, we drafted a set of both generic and domain-specific guidelines. These rules make the annotation process exhaustive and consistent, hence ensuring high data quality. The generic guidelines comprise the following commonly accepted best practices for object detection:

1. Draw a tight bounding box (BB) around an object of interest. Objects of interest must be (reasonably) sharp.

2. *Annotate every object of interest*. Objects of interest that are present in images but not annotated ‘confuse’ AI models (they are considered background).

3. *Extend BBs of occluded objects*. Sometimes an object is partially covered by another object in an image. If that is the case, ensure that the occluded object is annotated with a BB as if it were in full view.

4. Annotate cut-off objects at image borders if at least half of an object of interest is visible.

Besides generic ones, domain-specific annotation guidelines are needed to cope with the difficulties shown in Fig 1. EWD is challenging since multiple species exhibit large visual similarities and because the set of images covers a variety of flower stages – roughly: in bud, blooming, fruiting stage – drastically impacting the appearance of the wildflowers. Therefore, we propose the following three additional rules:

5. *Use floral count units as entities for annotation*. For most flowering plant species FCUs map one-to-one to botanically defined inflorescences. See also S1 Appendix.

6. *Only annotate wildflowers in blooming stage*. More precisely, for solitary flowers we propose to i) start annotating when flowers are open AND at least half of the petals are fresh, and ii) stop annotating when colours of the flower fade OR less than half of the petals are present. For the other FCUs, we propose to i) start annotating when at least one of the flowers belonging to the FCU is open and fresh, and ii) stop annotating when colours of the FCU fade OR blooming flowers are a minority of the FCU.

7. *Only annotate floral count units that are larger than 10mm* containing flowers that grow in easy-to-distinguish clusters. For solitary flowers ‘size’ refers to the dorsal-ventral axis. For other inflorescences it refers to the length of the largest axis. Motivation of this rule is that – given the striking visual similarities between closely related plant species – enough details of the FCU must be visible for proper identification by AI.

Fig 5 illustrates how these seven rules work out in practice. Strictly applying the above rules enable the AI to unlearn unwanted patterns, such as objects that are either out of scope (e.g., flowers that are in bud or withered) or of low quality (e.g., unidentified flowers that appears as unsharp coloured blobs); by doing so they are treated as background. Annotations are added with the open-source tool labelImg and are stored per image in PASCAL VOC format [21]. After the annotation process EWD consists of 2002 {<DATETIME-habitat>.jpg, <DATETIME- habitat>.xml} file pairs. The image file (*.jpg) and annotation file (*.xml) of each pair are given the same unique name that encodes date and time of the image exposure followed by one of the five landscape types.

**Fig 5:**
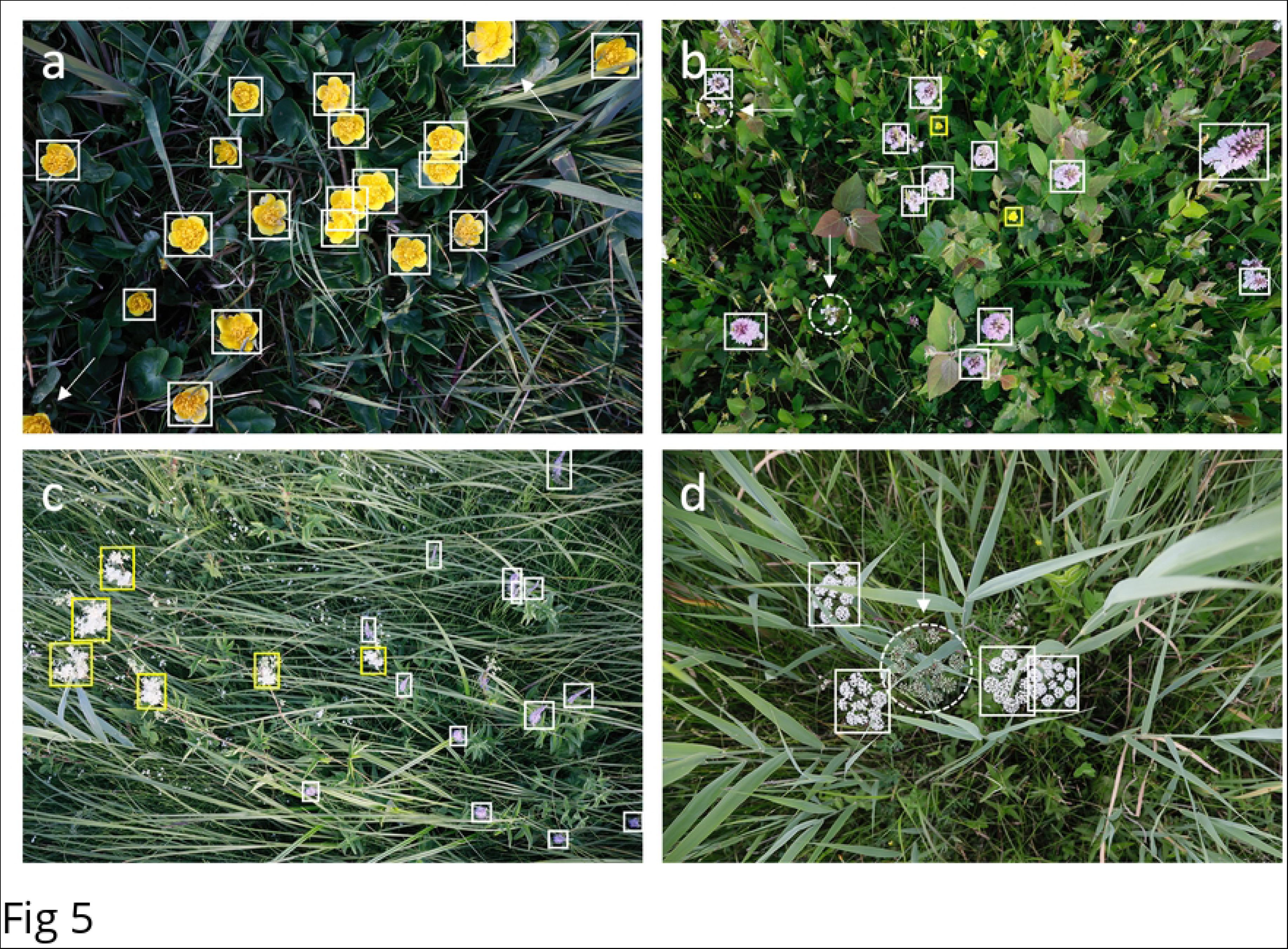
Annotated relevées compliant with the above guidelines. Note the tight and sometimes overlapping BBs (rule#1, rule#3). From top left to bottom right: a) *Caltha palustris,* the arrows point at cut-off flowers, they are only annotated when at least half of the FCU is visible (rule#4), b) *Dactylorhiza maculate* (white BBs), the arrows and dashed circles mark spikes that contain no open flowers (rule#6), and *Ranunculus repens* (yellow BBs) annotated as *Buttercup\**, c) *Veronica longifolia* (white BBs) and *Filipendula ulmaria* (yellow BBs), the ubiquitous tiny 2-3mm *Galium palustre* ‘speckles’ are not annotated (rule#7), d) *Cicuta virosa,* the arrow and dashed circle mark an umbel that is still in bud and hence not annotated (rule #6).

Summarizing, EWD holds 65571 annotations for 160 species and exhibits a long-tailed species distribution (which is typical for biological field data); 19 wildflowers have 1000 or more annotations per species, 22 wildflowers have annotation counts in the range [500, 999], 61 wildflowers have annotation counts in the range [100, 499], and 58 species have less than 100 annotations. The most common species in EWD is *Buttercup* (aggregate)* with 4196 annotations, followed by *Berteroa incana* with 3177 annotations. The number of annotations per image varies from 0 (some images have no annotations, e.g. because the depicted wildflowers are not in blooming stage) to a maximum 704 with an average of 32.75 annotations per image. The number of annotated species per image varies between 0 and 7, with a median of 1. Table 2 presents the EWD key figures per month.

**Table 2:** EWD key figures per month. The landscape types are abbreviated with R (roadside), U (urban green), C (cropland), G (grassland), and M (marsh).

| Blooming period | # images | # images per landscape type {R, U, C, G, M} | # species | # annotations |
| --- | --- | --- | --- | --- |
| **Year 2021** | **994** | **{419, 122, 42, 220, 191}** |  | **38515** |
| April | 75 | {18, 14, --, 38, 5} | 10 | 3103 |
| May | 296 | {69, 31, 12, 123, 61} | 46 | 14297 |
| June | 205 | {108, 18, 25, 13, 41} | 51 | 11585 |
| July | 215 | {63, 40, 5, 27, 80} | 56 | 4745 |
| August | 146 | {125, 19, --, --, 2} | 45 | 3421 |
| September | 57 | {36, --, --, 19, 2} | 20 | 1364 |
| **Year 2022** | **1008** | **{413, 115, 11, 237, 232}** |  | **27056** |
| March | 68 | {9, 9, --, --, 50} | 10 | 1765 |
| April | 122 | {19, 34, --, 66, 3} | 16 | 3983 |
| May | 209 | {99, 13, --, 63, 34} | 54 | 6661 |
| June | 298 | {102, 49, 11, 51, 85} | 75 | 7817 |
| July | 211 | {111, 10, --, 48, 42} | 60 | 4451 |
| August | 80 | {59, --, --, 5, 16} | 47 | 1767 |
| September | 20 | {14, --, --, 4, 2} | 16 | 612 |

Per species statistics of EWD, along with the defined FCU, the average number of individual flowers per FCU and the average number of FCUs per flowering plant, are given in S1 Appendix.

## 4. Data-centric AI demonstrator

To demonstrate the potential use of EWD, we trained an object detection model based on industry standard tools. More specifically, we used the open-source PyTorch framework and selected a state-of-the-art F-RCNN model [67], as provided in the TorchVision library. This end-to-end deep learning object detection model consists of a visual encoder (ResNet50) with pretrained weights followed by a classification and localization head that jointly learn in a multi-task setting [56, 68]. The choice for transfer learning with standard tools was motivated by the fact that we do not necessarily aim to train the best model possible but rather seek to explore the impact of improving the quality of the data on the model’s ability to estimate species richness and abundance. Thus, we opted for creating a baseline model with a relatively simple and fast training procedure and put more effort into preprocessing and balancing the data.

### 4.1. Preprocessing the data

Given as the EWD images have a resolution of 6720x4480 pixels and that the pretrained model has a maximum input size that is significantly lower (1333 pixels on either axis), the default behaviour, automatically resizing the image to match the maximum input size, drastically reduces the resolution of the EWD data. This might inhibit the model’s ability to correctly identify detected wildflowers. Our solution to this problem is to use a mosaic of the original EWD images by cutting them in tiles that are (roughly) equal to the maximum input size, eliminating or strongly reducing the degradation of image resolution. However, as shown in Fig 6, a simple cutting approach would inadvertently cut through annotated FCUs, which would make them less suitable as training data. Therefore, we devised a cutting approach that is aware of the presence of annotations in the image and attempts to find a less destructive way to cut the image into a mosaic. This produces a trade-off situation between perfectly matching the maximum input size of the F-RCNN model on the one hand and cutting through none of the annotations on the other hand. Our approach uses a 20% margin on the model’s maximum input size and then finds the cut lines that are least destructive within that margin.

**Fig 6:**
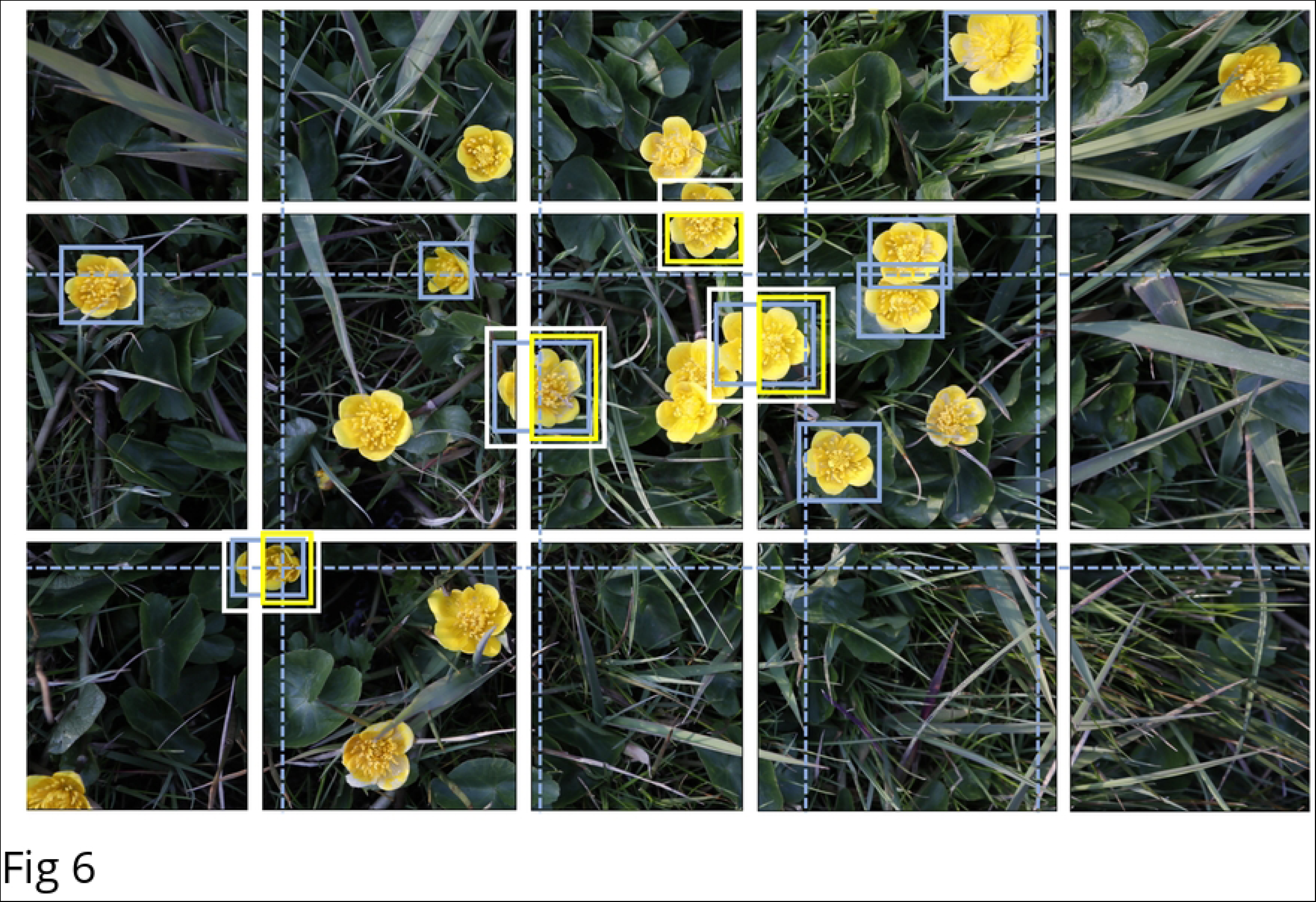
For the mosaic of the 30.4 Mpixel EWD images a setting was chosen that results in 15 (5×3) tiles. Starting with equally sized tiles (blue dashed lines) margin modifications of +/- 20% are allowed to minimize object cuts. In this example the number of object cuts is reduced from nine (blue BBs) to four (white BBs). To prevent double counts only the largest part of cut-off objects (yellow BBs) – in accordance with rule #4 of Section 3.3 – is added to the set of tiled annotations. White and yellow BB are drawn a bit larger for cosmetic reasons.

### 4.2 Balancing the data

The next issue to address is the highly imbalanced nature of the EWD (long-tail dataset), having some wildflowers being overly abundant and others being rather rare. Which, when used as training data, is likely to create bias in the model because of the classifier having insufficient training opportunity on sparsely represented classes and will probably favour overrepresented species [69]. Again, we choose a solution that influences the input data rather than manipulating the training process [70], using over/under-sampling techniques, or even adopting complex learning solutions such as for example adversarial neural networks that can potentially correct for bias to some degree [71, 72].

The solution we propose is to train with a balanced subset of EWD, i.e. we use a fixed predefined number of annotations per species for training, evaluating, and testing. To create this subset, we subsequently go through the tiled images of EWD selecting a combination of tiles that produces the required numbers. A similar approach is also adopted by Koch et al. [73] for bird images extracted from a Norwegian citizen science platform. For our object detection case this is not a straightforward process as while selecting an image tile containing annotations for one specific species, it may collaterally include annotations of other species as well. Although it is theoretically possible to calculate all tile permutations of possible solutions that bring to the exact number of required annotations per species maximizing the total number of species included, this is in practice not feasible due to its extreme high computational load. Hence, we followed a near-ideal heuristic approach. This approach guarantees that the preset number of required annotations per species are selected but as a drawback may drop a flowering plant species when a solution is not found within a certain number of attempts. Following this approach, we generated a stratified partition for 49 flowering plant species with the following fixed number of annotations per species: 250 for training, 50 for evaluation, and 50 for testing. Note that these numbers are not hard-coded but parameterized, see S2 Notebook.

### 4.3 Training the model

A relatively simple and fast training process with standard data augmentation practices using transfer learning is implemented. Training happens in train-and-evaluate loops, called epochs. Instead of fixing the number of epochs, after every epoch the evaluation metric ‘mAP @IoU > 0.5’ is compared to the average of this metric for the last five epochs, and if the value for the current epoch is lower than this average, the training process is stopped. In practice, the backpropagation process needs 13 to 15 epochs to converge, see S2 Notebook.

### 4.4 Performance of the model

The optimized model after training is able to generate predictions consisting of BB proposals with species labels and confidence scores. The overall model performance (Section 2.4) on the holdout test set has a mAP of 0.82 (@IoU > 0.5).

The AP results for each of the flowering plant species in the test set are shown in Fig 7. For the first 48 species the AP score gradually decreases from 0.99 for *Centaurea cyanus* to 0.45 for *Ranunculus aquatilis*. The rightmost species *Anthriscus sylvestris* is difficult to detect and shows a sharp drop to an AP of 0.19. The AP score is weakly correlated with the complexity of the species FCU. Most high score species have simple FCUs (solitary flowers, spikes, capitula), whereas the majority of wildflowers with low scores have complex FCUs (like racemes, cymes, corymbs and umbels).

**Fig 7:**
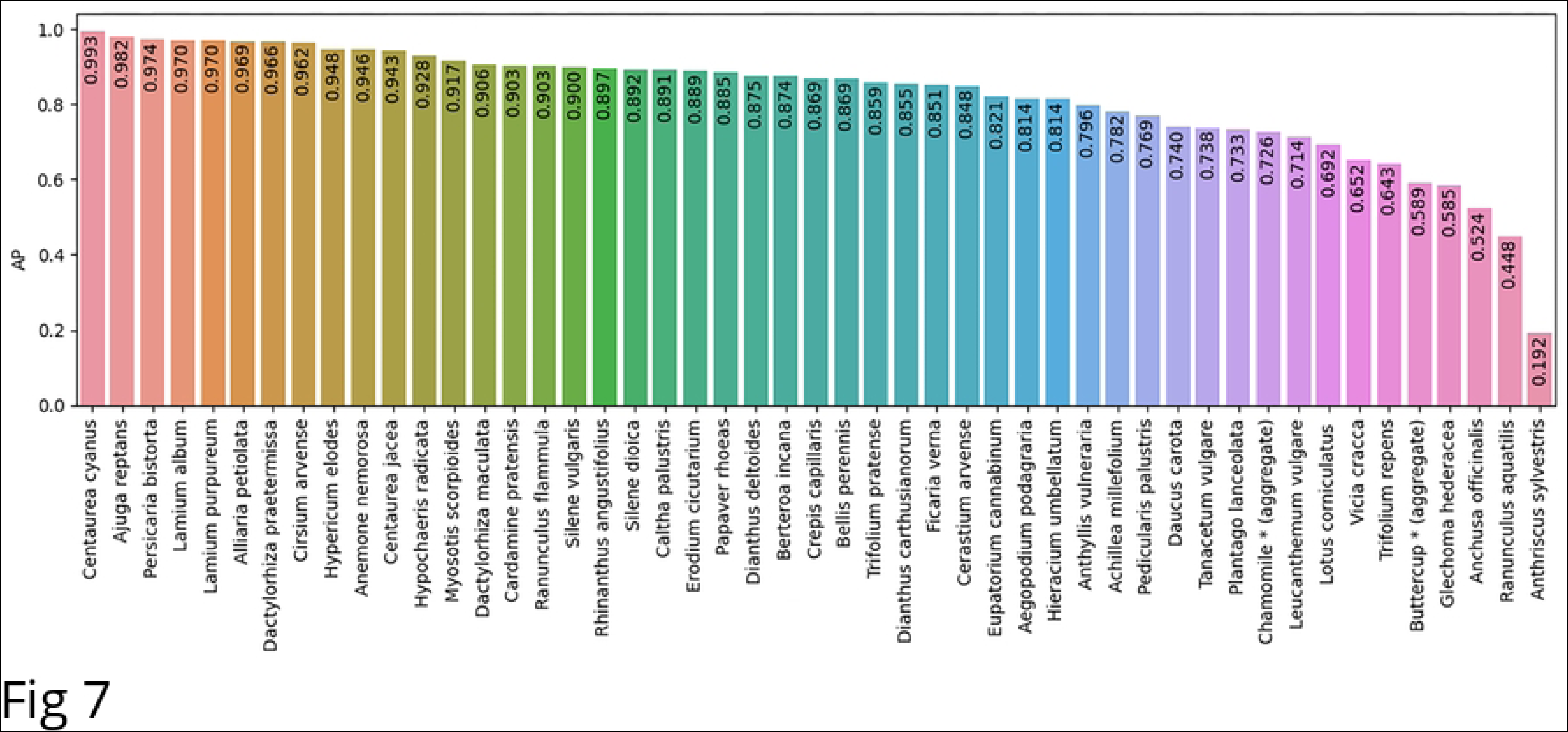
Average precision (AP) for each the 49 flowering plants.

More insight into counts and miscounts of the model is provided by the confusion matrix. Fig 8 shows the confusion matrix for the test set with detections that have a confidence score > 0.5. The rows of the matrix hold the actual annotations (they add up to 50 for each species); the columns indicate predictions. A perfect object detection model results in a confusion matrix with no off-diagonal counts.

**Fig 8:**
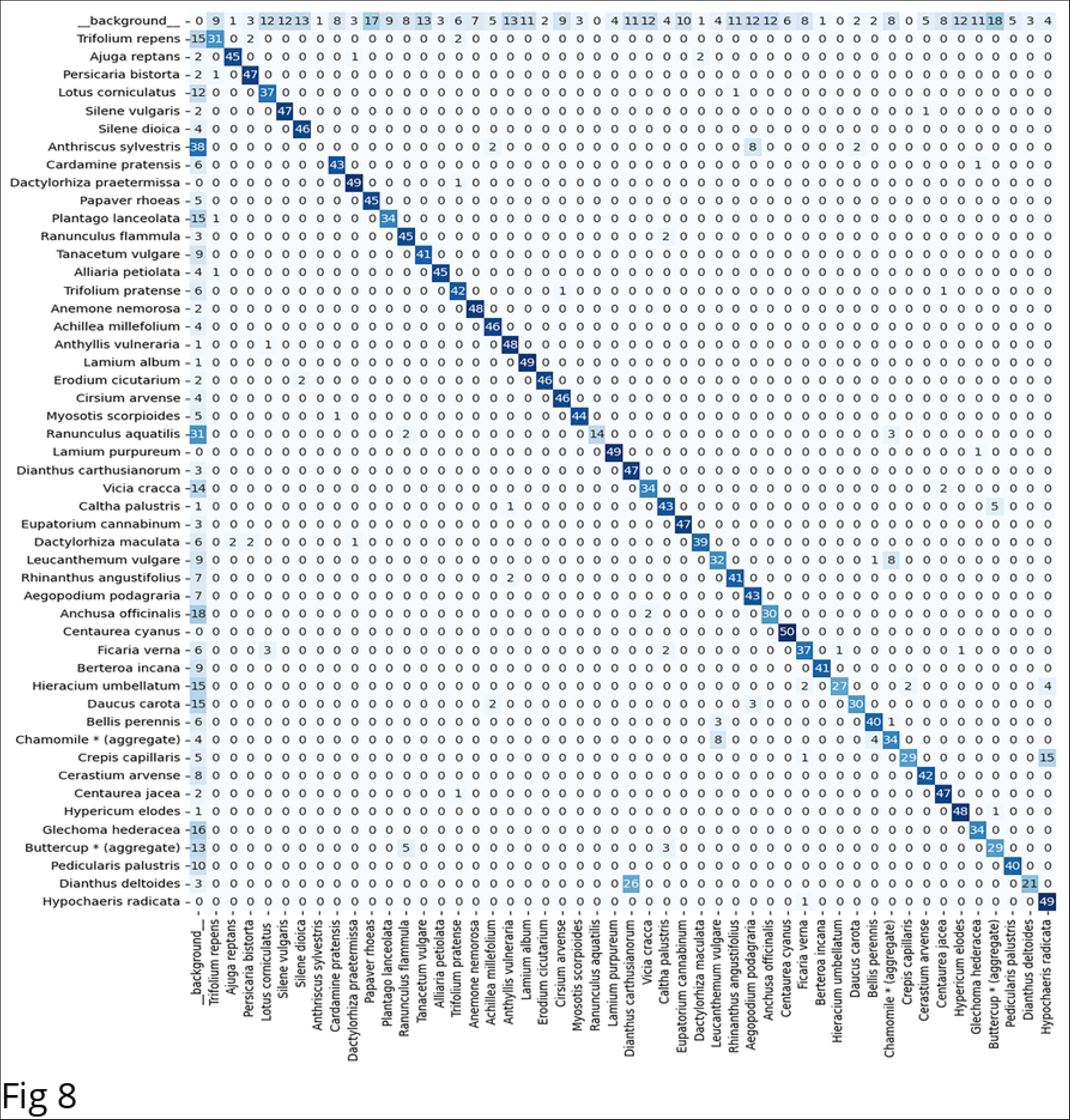
Confusion matrix for the holdout test set. The rows hold true wildflower annotations, the columns contain detections generated by the baseline model. The row labelled _background_ refers to spurious detections; the column labelled _background_ holds undetected species. The confidence score threshold used to construct this confusion matrix is set at 0.5.

Note that the diagonal is by far the most conspicuous pattern in the confusion matrix, indicating that the model generalizes well and is able to come up with reasonable wildflower counts retrieved from unseen test images. However, there are also a few other noticeable patterns. First of all, the row labelled as _background_ shows that there are quite some spurious wildflower detections in the background parts of the image, and secondly, the column labelled as_background_ demonstrates that a fair number of annotated wildflowers still go undetected. Species that have particularly low diagonal scores are *Anthriscus sylvestris* (also lowest AP value)*, Ranunculus aquatilis,* and *Dianthus deltoides.* Many *Anthriscus sylvestris* annotations (38 out of 50) and *Ranunculus aquatilis* annotations (31 out of 50) go undetected because they have low confidence scores (below the threshold) and are therefore treated by the model as background. *Dianthes deltoides* counts suffer less from the abovementioned ‘disappearance-in-the-background’ problem (3 out of 50) but are mainly confused with *Dianthes carthusianorum* (26 out of 50), a species from the same family. On the other end of the spectrum, species that can be counted very well by the model are the eye-catching sky blue *Centaurea cyanus* (50 diagonal hits; also highest AP value), *Dactylorhiza praetermissa*, *Lamium album*, *Lamium purpureum* and *Hypochaeris radicata* (all 49 diagonal hits). Furthermore, a notorious confusion is found between the two look-alike yellow composite species *Hypochaeris radicata* and *Crepis capillaris*.

## 5. Discussion

This paper presents two data products: i) a reference dataset for wildflower monitoring, and ii) an object detection model that can identify and count flowering plants from top-view images.

The EWD dataset holds 2002 images with 65.6K expert annotations for 160 species, belonging to 31 different families and is collected in two entire flowering seasons (begin of March till end of September). We have substantially extended FCU definitions for species with more complex (nested) inflorescences and our annotation process is supported by strict guidelines. This makes EWD a reference dataset that can be used for benchmarking computer vision algorithms. Moreover, we have added complementary botanical information to convert detected FCUs to individual flower or whole plant counts, thereby supporting other use cases related to wildflower monitoring.

Our baseline model – trained on a balanced subset of EWD – enables automated wildflower monitoring for 49 species with a mAP of 0.82. This high score makes the knowledge encapsulated by model actionable, i.e., it can be applied in practice for automatic large-scale wildflower monitoring. Other F-RCNN object detection models for wildflower monitoring are published by Hicks et al. [39] and Gallman et al. [60]. However, these studies, having 25K and 10K annotations respectively, collected wildflower data in limited period of one flowering season (May till August) and their models are able to identify and count only 25 flowering plant species, half the number of species that our model can handle. In contrast to our study, they did not follow a strict annotation protocol and focused primarily on the model, not on the data perse. The model of Hicks et al. [39] is trained with simple inflorescences (only solitary flowers and capitula) and did not report the generally accepted object detection mAP metric. The Gallman et al. study [60] did not explicitly use FCUs and reported a mAP score of 0.48.

### 5.1. Quality of annotations

The first data product, EWD, is an expert-annotated dataset. Platforms like Amazon’s Mechanical Turk have made it possible to outsource annotation work as a crowdsourced task, at relatively low cost. Although several methods – either ex-ante or ex-post – exist to correct for unreliable labels of crowd workers [74–77], this remains a second-best approach [78]. High-quality consistent expert annotations along with a well-defined data acquisition protocol are paramount for a data-centric AI approach. They turn datasets into reference datasets. By providing a solid ground-truth, reference datasets allow for benchmark studies, varying from comparing different computer vision algorithms, developing metrics, to evaluating innovative techniques such as semi-supervised learning [79, 80] that have the potential to alleviate the burden of adding huge numbers of manual annotations.

Although the quality of the EWD annotations is even further enhanced by formulating explicit guidelines (Section 3.3), some rules are hard to objectify. Especially rule#1 could be made more explicit by exploring the use of state-of-the-art no-reference image quality metrics for sharpness [81].

### 5.2 Model limitations and possible improvements

The second data product is our wildflower object detection model. It builds upon a F-RCNN backbone; an end-to-end deep learning architecture that is a proven solution ‘from the zoo’. The model demonstrates the potential use of the EWD dataset. It can be adapted easily to accomodate more wildflowers by lowering the required number of annotations per species (currently set at 250 for training, 50 for evaluation, and 50 for testing). The main motivation of the modelling choices made is preserving data quality. A critical reflection on the modelling choices is given below.

Although our preprocessing solution – partition the image and feeding high resolution tiles to the F-RCNN – avoids the tenacious ‘small object detection’ problem [82], it creates a minor artifact as well. In particular, the smaller parts of cut-off objects (for which the annotations are removed, see Fig 6) generate quite some spurious detections, even though the model is trained with annotations that contain at least half of the FCU (see Section 3.3; rule #4). Another artifact arises from balancing the data. Because of this, the model is trained on a subset of flowering plant species and might easily generate false hits in those few test images that happen to hold look-alike species on which it was not trained. Both imperfections account for the relative high number of hits in the _background_ row of Fig 8.

Transfer learning is often perceived as inferior to full ‘training-from-scratch’ schemes. However, we view bypassing the vast number of weights that belong to visual encoders in the training process not as a limitation but as a less-is-more benefit. It not only keeps the model optimizing process fast and flexible but also strongly reduces the carbon footprint of the training the AI [83]. Moreover, recent literature [84] indicate that using pre-trained networks in many practical situations even lead to performance gains. Hence, we rather keep the model optimizing process fast and flexible, i.e. stick to the transfer learning paradigm, and explore in our future work: i) cost-function engineering tailored to long-tailed distributions, such as proposed by Tan et al. [85], and ii) more fundamental ‘beyond vision’ solutions as put forward below.

The most important shortcoming of object detection models is that proposed bounding boxes with a class and score are considered as independent entities and that predictions are only based on local visual content. They have no mechanism to explicitly relate localized predictions in the same image to one another. In the physical world, visual objects usually coexist with other related objects (a fortiori for wildflowers), human perception very much relies on this fact [86]. Henceforth, an intelligent agent or voting procedure that considers neighbouring wildflowers might boost model performance significantly. To give a trivial example: the best species predictor for a non-canonical view of an occluded flower that is hard to recognize is probably a look-alike species (same colour) in the proximity. So, picking matches from neighbouring species labels for predictions with a low confidence score might be a promising direction to explore. Another limitation is that our wildflower monitoring framework is agnostic with respect to flowering time and landscape type. Ecologists implicitly include this type of information while doing fieldwork, i.e., they implicitly rule out or confirm – based on date and habitat – ‘suggestions’ created by their visual system. Modifying confidence scores of wildflowers by explicitly encoding this type of domain knowledge is envisioned as another promising future direction.

### 5.3 Ongoing work and outlook

Current work focuses on extending the ‘AI demonstrator’ (Section 4) with: i) a user interface supporting interaction with different stakeholders [87], and ii) a mobile sensor with guidance for correct image acquisition and streaming cloud storage. These mobile sensors could be drones for professional use or smartphones for the general public.

In future work the presented method for automated wildflower monitoring will be validated in the field and extended to other flowering plant species. It is a valuable tool that can be used for ecological and environmental research, e.g., to investigate whether wildflower occurrence and richness can be used as a proxy for mowing regimes, nitrogen depletion or climate change. Policy makers might use the results of automated wildflower monitoring for spatial planning activities and biodiversity conservation [88]. In the long run, we envisage that AI-enabled monitoring solutions will be tied to more coarse-grained ecological indicators [89].

Finally, we hope to motivate and engage AI researchers to use the EWD reference dataset and create improved solutions for automatic wildflower monitoring.

## 6. Conclusions

Humans have a knotty relation with nature. On the one hand, using technology we put a tangible mark on our planet causing climate change and an unprecedented biodiversity loss. On the other hand, our technology has the potential to shape transitions towards a more sustainable world. This study belongs to the latter category. It aims at showcasing automated wildflower monitoring using AI. Wildflower diversity is a profound part of our natural capital.

More specifically, this paper demonstrates how the quality of automated wildflower monitoring can be enhanced by adopting a data-centric AI approach. Distinct aspects of our work that treat data as a first-class citizen are: i) a protocol for collecting relevées (Section 3.1), ii) the introduction of FCUs for a wide variety of flowering plant species (Section 3.2), iii) expert annotations according to strict guidelines (Section 3.3), iv) preprocessing that preserves image resolution (Section 4.1), and v) training the F-RCNN-ResNet50 object detection model with a balanced subset in order to obtain unbiased predictions (Section 4.2).

This baseline model can recognize and count 49 lowering plant species with a mAP of 0.82, thereby outperforming contemporary automated wildflower monitoring solutions. We hope to inspire and encourage the computer vision research community to create improved AI-enabled solutions for wildflower monitoring using EWD as a reference dataset.

## Supporting information

S1 Appendix: EWD species characteristics and statistics.

S2 Notebook: Python code to train and test the object detection model.

## Data availability statement

EWD is publicly available under the CC BY-NC 4.0 license for non-commercial usage on the DataverseNL research data repository at https://doi.org/10.34894/U4VQJ6.

